# Sensorimotor mapping of volitional facial movements in Tourette syndrome

**DOI:** 10.64898/2026.04.02.712172

**Authors:** Caitlin M. Smith, Mairi S. Houlgreave, Michael Asghar, Susan T. Francis, Stephen R. Jackson

**Affiliations:** School of Psychology, University of Nottingham; Precision Imaging Beacon of Excellence, School of Medicine, University of Nottingham; Centre for Neurotechnology, Neuromodulation and Neurotherapeutics, School of Psychology, University of Nottingham; Sir Peter Mansfield Imaging Centre, School of Physics and Astronomy, University of Nottingham

**Keywords:** Tourette Syndrome, tics, facial tics, volitional movements, sensorimotor, functional representations, functional magnetic resonance imaging

## Abstract

**Background:** Tourette Syndrome (TS) is a neurodevelopmental movement disorder involving involuntary motor and vocal tics believed to be characterised by disordered neural inhibition. Cortical representations have previously been manipulated by disruptions in the inhibitory neurotransmitter γ-aminobutyric acid (GABA). However, while facial tics are the most reported motor tic, it is unclear if facial sensorimotor representations differ in TS.

**Methods:** Sixteen individuals with Tourette Syndrome (TS) or chronic tic disorder and twenty typically developing (TD) control participants underwent 3-Tesla functional magnetic resonance imaging (fMRI). Blood-oxygenation level-dependent (BOLD) responses were measured during a block-design task comprising cued facial movements of common facial tics (blinking, grimacing and jaw clenching). Activations in bilateral pre- and post-central cortices and supplementary motor areas (SMA) were examined. Conjunction analyses identified voxels commonly and uniquely activated across movements within each group.

**Results:** Both groups showed significant activations in the bilateral sensorimotor cortices and SMA in response to blink, grimace and jaw clench movements, with no significant between-group differences. Between-group similarities were lowest for unique blink maps. Common voxel maps also revealed low between-group similarity, with reduced sensorimotor activation and no shared SMA activation across movements in the TS group.

**Conclusion:** Voluntary facial sensorimotor representations do not differ between groups. However, low similarities between group unique blink maps may reflect greater prevalence of blinking tics in TS. Additionally, reduced overlap in sensorimotor activation and absent common SMA engagement across cued movements in the TS group may indicate altered motor integration or action initiation.

## Introduction

Tourette Syndrome (TS) is a neurodevelopmental hyperkinetic movement disorder, characterised by the repeated occurrence of involuntary and stereotyped motor and phonic tics [1]. Facial tics (such as blinking, grimacing and jaw clenching) are the most commonly reported motor tic and are often the first to appear [2–5]. TS is thought to originate from aberrant activation within cortico-striatal-thalamo-cortical (CSTC) circuitry due to striatal disinhibition [6–12] and is associated with abnormalities in sensorimotor inhibitory functioning [13–19].

While tic phenomenology greatly varies between individuals, specific tic presentation has been theorised to stem from aberrant activity and disinhibition within discrete regions of topographically organised basal ganglia nuclei. This has been demonstrated in rodent models of TS, whereby delivering injections of *y*-aminobutyric acid A receptor (GABA_A_) antagonists to focal somatotopic areas of the striatum resulted in tic-like movements in specific body parts [7]. However, it is believed that facial muscles may be preferentially recruited in humans with TS due to stronger representation and cortical magnification of facial regions within topographical maps across the primate CSTC circuit [20–22]. This is supported by non-human primate models, where striatal disinhibition results in orofacial tic-like movements, regardless of the topographical (GABA_A_) antagonist injection site [6, 23–26]. This might therefore explain the commonality and initial appearance of facial tics in TS.

GABA_A_ is also implicated in the mediation of sensorimotor representations due to its proposed role in implementing surround inhibition [27–31]. However, GABA functioning alone may not be sufficient to produce changes in sensorimotor representations. For instance, Focal Hand Dystonia (FHD) is characterised by involuntary movements and spasms of the hand and is associated with bilateral abnormalities in sensorimotor inhibition [32–33]. However, abnormalities in cortical representations are limited to the affected digits and associated hemisphere [34], with bilateral abnormalities associated with greater symptom duration and severity of condition [35–36]. As a result, altered representations are proposed to occur due to dysfunctional cortical inhibitory mechanisms, paired with repetitive behaviours of the hand [37–38]. It is proposed therefore that the presence of altered sensorimotor inhibition and repetitive facial tics in TS may be sufficient to encourage malleability of facial cortical representations.

This study used 3-Tesla functional Magnetic Resonance Imaging (fMRI) to investigate sensorimotor cortical representations during volitional facial movements relating to common tics (blinking, grimacing and jaw clenching). We hypothesised that there would be differences in the sensorimotor representations of facial movements between individuals with TS and typically developing (TD) controls.

## Methods

### Participants

Twenty TD controls and twenty participants with TS or chronic tic disorder took part in the study. Four participants in the TS group were excluded from subsequent analysis due to excessive head motion during fMRI, with the resulting demographics in each group described in Table 1. Age (*U* = 127, *p* = 0.292) and sex (X^2^ (1, *N =* 36) = 0.00, *p* = 1.00) distributions were not significantly different between groups.

**Table 1.** Demographic information of TD and TS participants.

| Group | N | Age (years) | Sex |
| --- | --- | --- | --- |
| TD | 20 | 31.20 ($\pm 9.8$) | 10 male, 10 female |
| TS | 16 | 35.06 ($\pm 12.2$) | 8 male, 8 female |

*Note.* N = number of participants, TD = typically developing, TS = Tourette Syndrome, mean ( ± standard deviation)

Participants gave informed consent, and study ethics were approved by the local ethics committee (School of Psychology, University of Nottingham: S1454). No participants in the TD group were taking central nervous system (CNS)-active drugs and all were free from any mood or neurological disorder. For safety and MRI data quality, individuals with TS were screened based on their tic frequencies/severities and only those without frequent and severe head tics were recruited. All participants with TS reported experiencing facial tics. Yale Global Tic Severity Scale (YGTSS) [39] and Premonitory Urge for Tic Disorders Scale – Revised (PUTS-R) [40] were administered to individuals with TS, with medication status, YGTSS and PUTS-R scores for the TS group presented in Supplementary Information 1. All participants were free of MRI contraindications and right-handed, as assessed by the Edinburgh handedness questionnaire [41].

### fMRI Paradigm

Three 4.5-minute fMRI task blocks were acquired, during which participants were visually instructed to perform facial movements at 1Hz for 8 s, followed by 24 s rest (repeated for 8 cycles). Stimuli were presented using PsychoPy (Open Science Tools, Ltd) [42] on a MR-compatible screen (BOLDscreen 32, Cambridge Research Systems, UK), viewed via a mirror mounted to the head coil. Each fMRI task block consisted of a single facial movement presented in a fixed order (blinking, grimacing, jaw clenching), all facial movements that are very common tics in TS [2–5]. Participants were trained on each facial movement prior to scanning to ensure consistency.

Video recordings were acquired during scans using a MR compatible camera with integrated LED light mounted to the head coil (12M-I, MRC systems GmbH). This was used to assess task compliance and to extract precise facial movement onset and duration timings to enter for inclusion as regressors in GLM analyses. Recordings were unavailable for all blocks for one TS participant and six TD controls due to lack of consent or technical issues; in these cases, fixed block timings were used.

### MRI data acquisition

MRI data were acquired on a Philips 3T Ingenia MRI scanner (Philips Healthcare, Best, The Netherlands) using a 32-channel head coil at the Sir Peter Mansfield Imaging Centre, University of Nottingham, UK. Three BOLD fMRI data sets were acquired using a single-shot 2D T_2_*-weighted gradient-echo (GE)-echo planar imaging (EPI) sequence (2.5 mm isotropic voxels, matrix 84 x 83 x 48, multiband 4, SENSE 1.5, TR/TE = 1000/30 ms, 4 min 16 s per block). Two additional spin-echo (SE)-EPI short scans with opposing fat-shift directions (5 dynamics; 2.5mm isotropic voxels, matrix 84 x 83 x 48, multiband 4, SENSE 1.5, TR/TE = 4000/60 ms) were acquired for distortion correction.

A T_1_-weighted 3D Magnetisation Prepared-Rapid Gradient Echo (MPRAGE) scan was acquired following fMRI (matrix size = 256 x 256, FOV = 176 x 256 x 256 mm, 1 mm isotropic, 176 slices, TR/TE = 8.1/3.7 ms, flip angle = 8 °, acquisition time 3 minutes 35 seconds). This was co-registered to the fMRI data and normalized to Montreal Neurological Institute (MNI) space for group-level comparisons.

### BOLD-fMRI analysis

#### Pre-processing

fMRI data were distortion corrected using FSL-TOPUP and denoised using Noise Reduction with Distribution Corrected (NORDIC) principal component analysis (PCA) [43–44]. Data were assessed for motion, and any fMRI blocks with absolute mean motion displacement >2.5mm were removed. To maximise data retention, especially in the TS group, blocks with brief motion contaminated segments (>2.5 mm, up to three cycles) were removed from subsequent analyses using FSL functions [45]. This resulted in exclusion of data from three TS from ‘blink’ blocks, one TD and eight TS from ‘grimace’ blocks and two TD and six TS from ‘jaw’ blocks.

Temporal SNR (tSNR) of de-noised fMRI timeseries were assessed in MATLAB (MATLAB_R2019a, Mathworks, Natick, MA) using in-house quality assurance code (https://github.com/nottingham-neuroimaging/qa/tree/master/fMRI_report_app). fMRI blocks with tSNR <30 were to be excluded from analyses; however, all fMRI blocks had tSNR above this threshold. Final participants numbers and mean tSNR (± SD) for each movement block are reported in Table 2.

**Table 2.** Number of participants and tSNR in blink, grimace and jaw clench blocks for TD and TS groups.

| Group | Blink |  | Grimace |  | Jaw clench |  |
| --- | --- | --- | --- | --- | --- | --- |
|  | N | tSNR | N | tSNR | N | tSNR |
| TD | 20 | 134 ($\pm 11$) | 19 | 115 ($\pm 8$) | 18 | 113 ($\pm 13$) |
| TS | 13 | 98 ($\pm 14$) | 8 | 99 ($\pm 20$) | 10 | 96 ($\pm 17$) |

*Note.* N = number of participants, tSNR = temporal signal-to-noise ratio, TD = typically
developing, TS = Tourette Syndrome, mean ( ± standard deviation)

Motion correction was performed using FMRIB’s linear image registration tool, MCFLIRT, using the middle volume as a reference volume [46]. Denoised data were pre-processed using FSL-FEAT (FMRI Expert Analysis Tool) [45], including high-pass temporal filtering (32 s) and spatial smoothing with SUSAN (3.75 mm FWHM) [47].

fMRI images were moved to MNI space. An example volume functional volume was first aligned FLIRT (FMRIB) [46,48] to the participant’s T_1_-weighted anatomical image which was normalised to MNI space. The combined transformations were applied to the fMRI images to move to MNI space for group-level analysis.

#### Standard imaging analysis

Single-subject and group-level analyses were conducted using FSL-FEAT. Standard and extended motion parameters were included, along with an additional confound regressor to scrub volumes with framewise displacement > 0.9 mm. Pre-whitening was applied to account for temporal autocorrelation.

For each fMRI scan, GLM regressors of interest were formed from the onset and duration of each facial movement (blink, grimace, jaw clench) derived from the video recordings (or block timings when unavailable) and convolved with a double-gamma haemodynamic response function. Movement-specific contrasts were defined for each condition (blink, grimace, jaw clench).

Group-level analyses were performed using mixed-effects modelling (FLAME 1+2) to estimate mean responses for the TS and TD groups for each facial movement. Results were masked to the bilateral supplementary motor area (SMA), and bilateral pre- and post-central cortex (Harvard-Oxford Cortical Structural atlas) and cluster-corrected (Z > 2.3, p = 0.05) [49]. Brain regions associated with Z-max MNI coordinates were determined in MATLAB R2019a using mni2atlas (https://github.com/dmascali/mni2atlas/releases/tag/1.1). The SMA was included in this ROI mask due to its role in action initiation and control and its reported dysfunction in TS [50–51].

#### Conjunction analysis

A conjunction analysis was performed in AFNI [52–53] to identify voxels commonly activated across thresholded group-level contrasts (blink, grimace and jaw clench) from FSL-FEAT analyses. Unique activation maps were generated by subtracting the conjunction map from movement-specific contrast. Dice coefficients were calculated to quantify the overlap of common and unique voxel maps for each facial movement between TD and TS groups.

## Results

### Cluster-level inferences

Significant cluster activations were identified in bilateral sensorimotor cortices and the SMA for blink, grimace, and jaw clench in both the TD and TS groups (Figure 1; Z = 2.3, *p* < 0.05), with no significant between group differences. Cluster size, Z-max coordinates, and the associated brain region for each cluster activation are reported in Supplementary Information 2.

**Figure 1.**
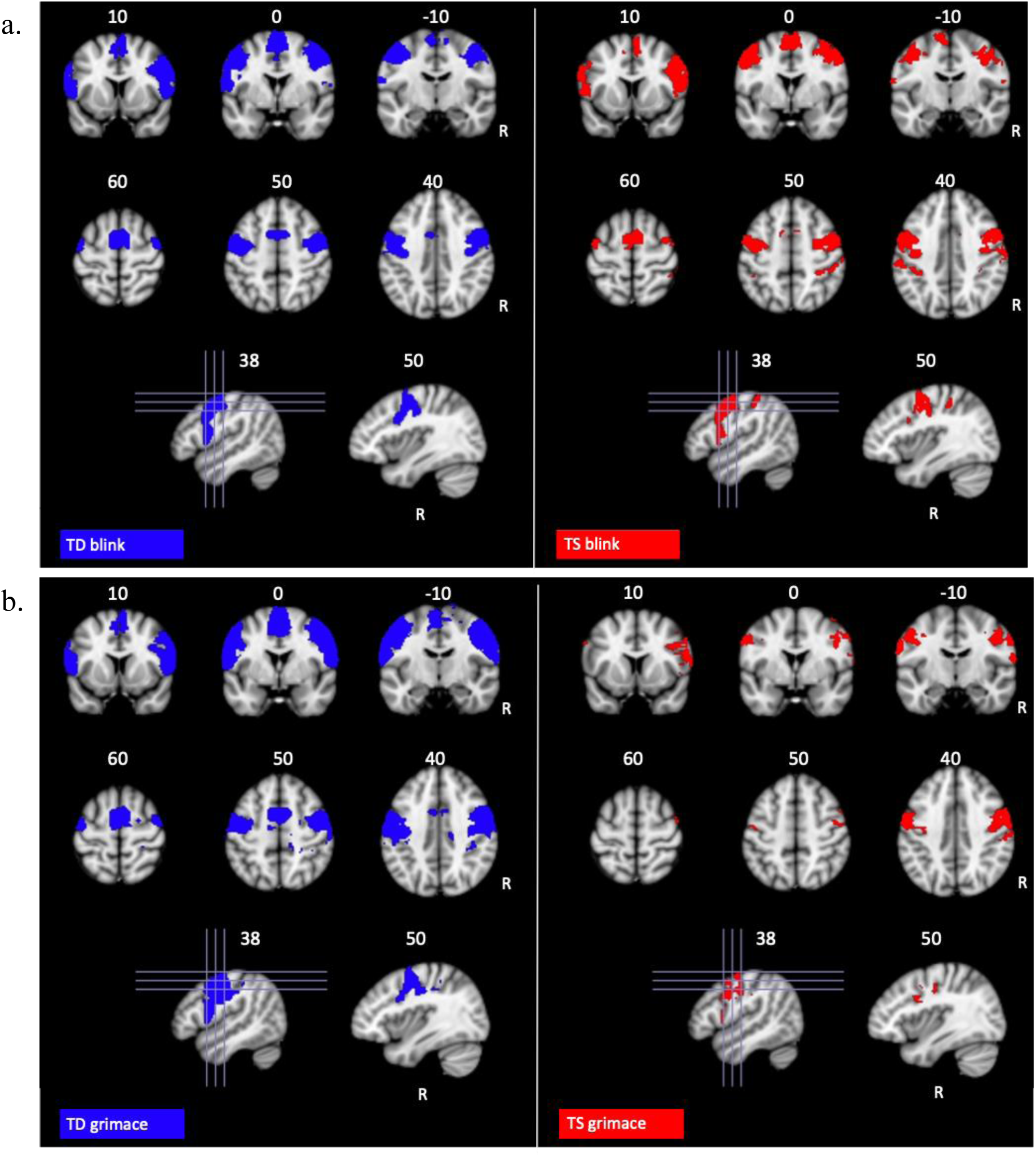
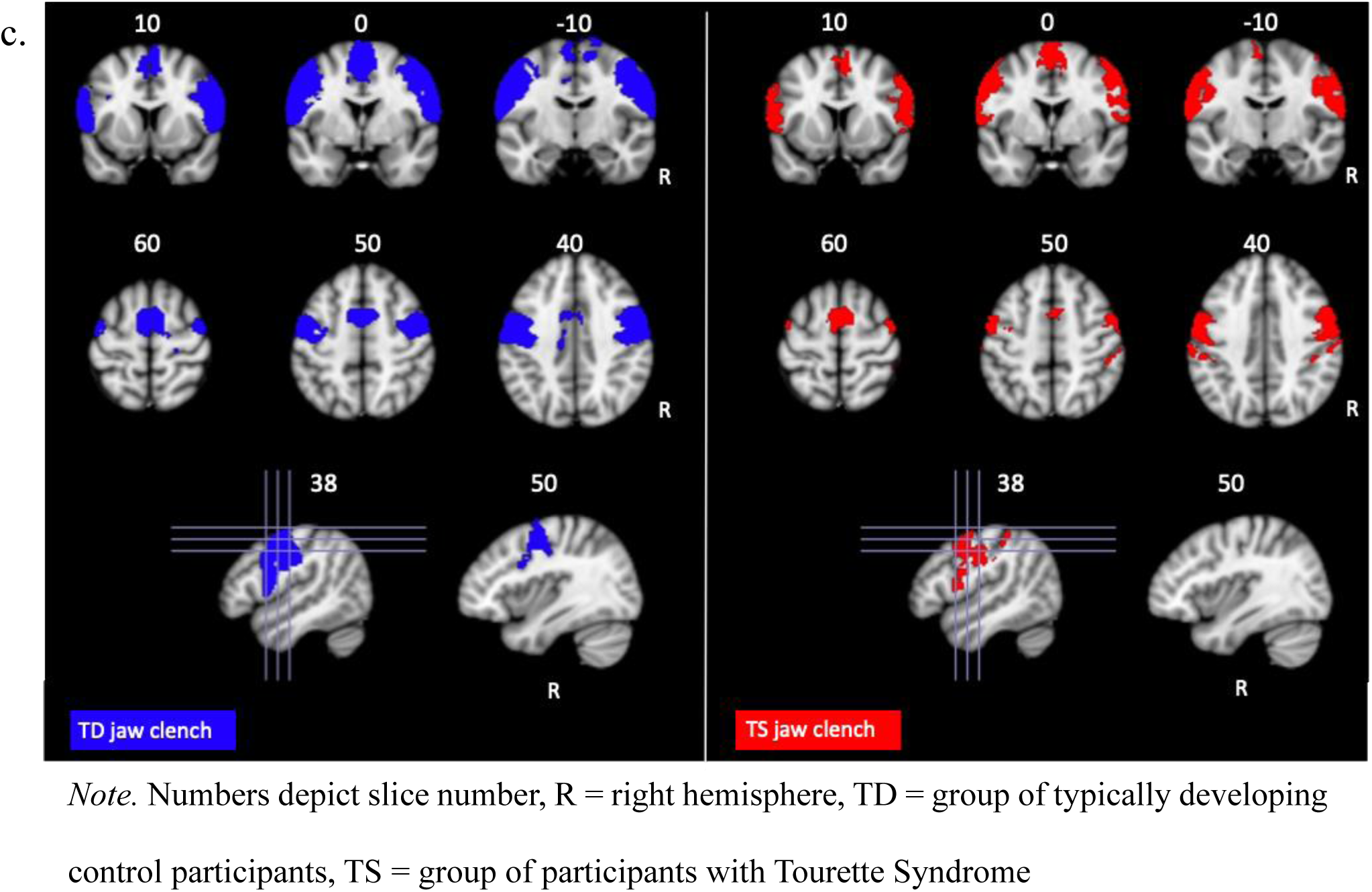
Cluster activations in TD (blue) and TS (red) groups for (a) Blink, (b) Grimace, and (c) Jaw clench blocks Note. Numbers depict slice number, R = right hemisphere, TD = group of typically developing control participants, TS = group of participants with Tourette Syndrome

### Conjunction analysis

Conjunction analysis identified unique (Figure 2) and common (Figure 3) voxel maps across all facial movements in both the TD and TS groups. The similarity of unique and common voxel maps between TD and TS groups is quantified using Dice coefficients in Table 3, revealing low overlap for common voxels. For unique maps, similarity was lowest for blink, followed by grimace and highest for jaw clench which showed moderate similarity.

**Figure 2.**
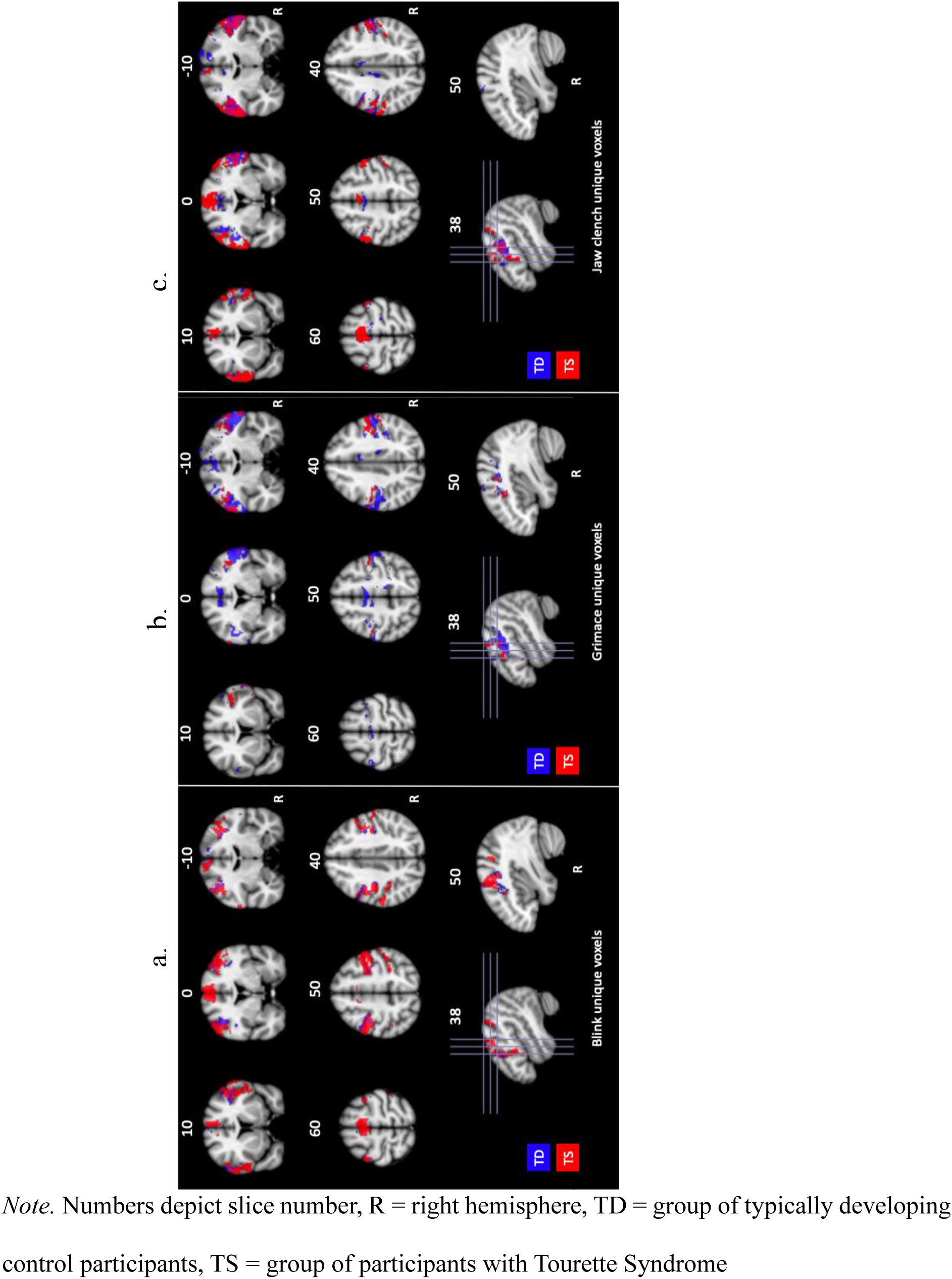
Unique voxels identified in a conjunction analysis in the TD (blue) and TS (red) groups to (a) blink, (b) grimace, and (c) jaw clench movements. Note. Numbers depict slice number, R = right hemisphere, TD = group of typically developing control participants, TS = group of participants with Tourette Syndrome

**Figure 3.**
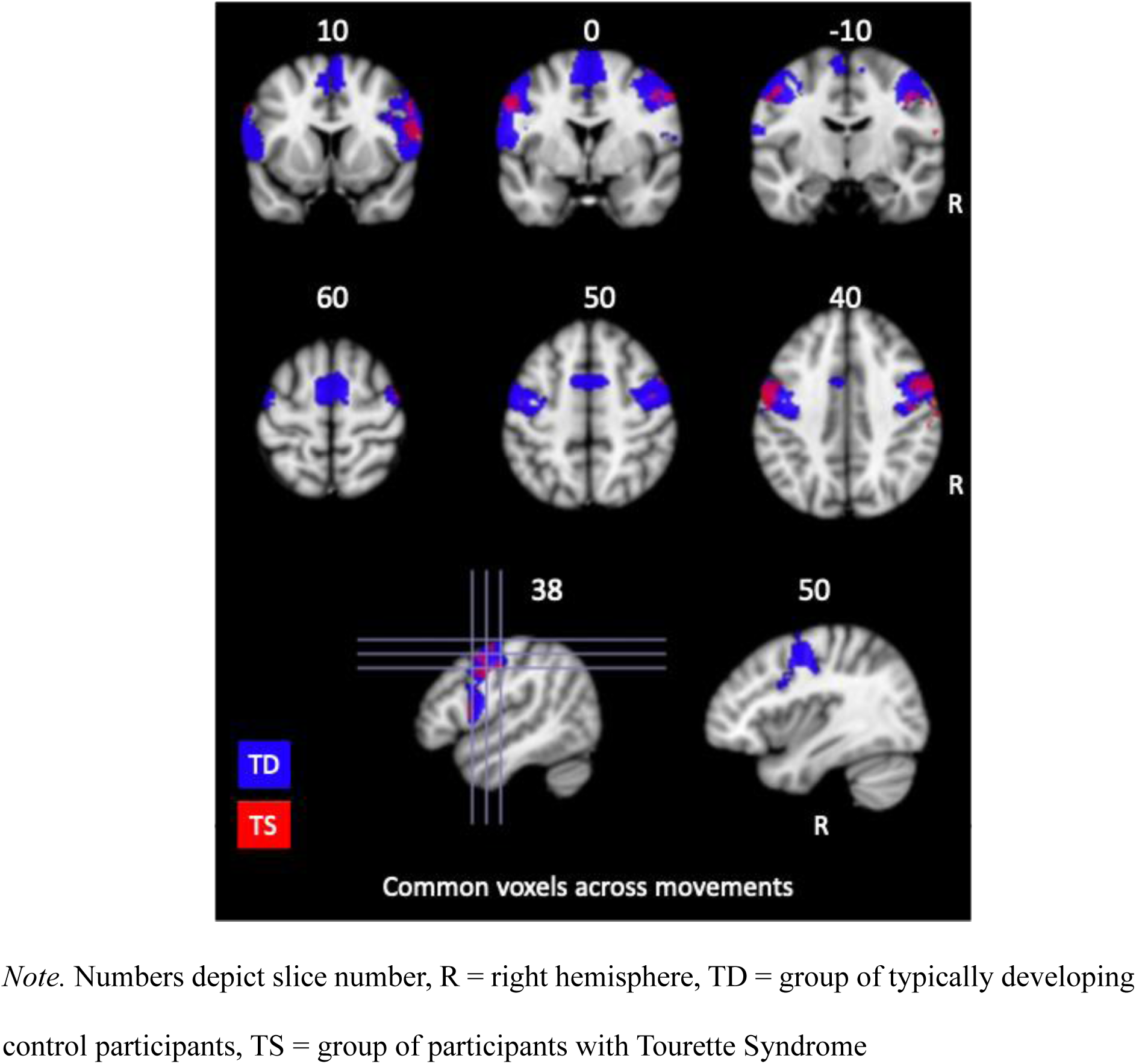
Common voxels identified in a conjunction analysis in TD (blue) and TS (red) groups across movements (blink, grimace and jaw clench) Note. Numbers depict slice number, R = right hemisphere, TD = group of typically developing control participants, TS = group of participants with Tourette Syndrome

**Table 3.** Dice coefficient between voxels unique to blink, grimace and jaw clench blocks and voxels common across all movement blocks between TD and TS groups.

|  | Unique |  |  | Common |
| --- | --- | --- | --- | --- |
|  | Blink | Grimace | Jaw Clench |  |
| Dice coefficient | 0.15 | 0.26 | 0.47 | 0.21 |

Visual inspection showed that unique maps for each movement exhibited overlap with the sensorimotor face region in both groups (Figure 2). Common voxel maps also overlapped the sensorimotor face region. However, while the TD group showed common activation within the SMA, no such overlap was observed in the TS group (Figure 3).

## Discussion

This study examined whether sensorimotor fMRI activations during facial movements mimicking common tics differ between individuals with TS and TD controls. Activations for each facial movement (blink, grimace and jaw clench) were observed across a sensorimotor mask encompassing bilateral S1, M1 and SMA. However, no between-group differences in cluster activation were identified for any movement, suggesting that cued tic-like voluntary facial movements engage similar sensorimotor and SMA responses in TS and TD groups. This contrasts with predictions of altered representations in TS due to impaired inhibition and repetitive expression.

The absence of group differences may be due to participants with TS engaging in volitional movements rather than spontaneous tic movements. While there is a debate on whether tics are involuntary movements or voluntary responses to premonitory urge phenomena [54], neural activity prior to voluntary movements show similarities between individuals with tic disorders and neurotypical controls, while differing from tic-related activity [55–56]. In addition, electrophysiological recordings of centromedial thalamic nuclei have shown differential patterns between the onset of voluntary movements and tics [57]. This suggests that neural markers of voluntary facial movements may be preserved in individuals with TS.

Conjunction analysis revealed that both groups showed unique and common voxel maps within the sensorimotor cortex. However, Dice coefficients revealed low similarity between groups, particularly for blinking, followed by grimacing and jaw clenching. This may reflect differences in the sensorimotor cortices when producing movements related to more prevalent tics, such as eye blinking which have a reported lifetime prevalence of 91.5% of those with TS [4]. This is supported by recent evidence in a sample of 227 adults with TS, that eye blinking was the most reported tic (55%), followed by simple mouth movements (28%) and grimacing (26%) [5]. However, tic prevalence was not assessed in the current study.

Dice coefficients of low similarity in common voxel maps between TD and TS groups further suggests differential recruitment of sensorimotor regions across groups. The TD group showed more extensive and consistent overlap, including within the SMA, whereas the TS group showed reduced overlap and no common SMA activation. This aligns with previous evidence of disordered activity within the SMA during tasks involving cognitive control of movement [50]. Given the role of the SMA in action initiation and volitional control of action and its link to tic suppression in TS [51, 58], this may indicate altered motor integration or control processes.

Task-specific differences in the TS group were also present. While SMA activation was present for blinking and jaw clenching, it was absent for grimacing, a more complex movement involving more facial muscles and requiring greater movement coordination. Previous evidence has shown that reduced motor cortical excitability is evident during tasks requiring greater cognitive control in TS participants, with reduced motor cortical excitability thought to be a result of increased SMA GABA, as measured with Magnetic Resonance Spectroscopy (MRS) [15, 50, 59–62]. In TS, SMA GABA has been shown to inversely relate to SMA BOLD responses [13]. This may explain reduced SMA engagement during the more demanding grimacing task as a result of greater inhibition.

Finally, review of video data during motor blocks highlighted that some facial tics were present during motor blocks, which may also have induced aberrant activation in the SMA. Prior evidence has examined cross-correlation activations between tics in individuals with tic disorders and specific volitional ‘tic-like’ movements in age- and sex-matched TD controls [63]. Similar patterns of cross-correlation were identified in activations from the motor cortex to the rest of the brain, however broader cross-correlations were identified between the motor cortex to the SMA in the group with tic disorders only. This indicated the presence of greater SMA activation prior to and after tic execution compared to volitional movement [63]. As a result, the presence of tics is another possible cause of the aberrant activation in the SMA across tasks in the TS group, which is consistently activated across tasks in the TD group. As the number of tics and level of suppression during tasks were not quantified here, this cannot be concluded.

## Limitations

Several limitations should be noted within this study. First, differences in GABAergic inhibition between groups was not assessed, as measures indicative of GABA concentration such as MRS-GABA or paired-pulse transcranial magnetic stimulation (TMS) measures were not collected. Therefore, the specific role of GABAergic inhibition mechanisms in shaping sensorimotor maps remains unclear.

Secondly, differences in sensorimotor activations to each facial movement may reflect individual symptom phenomenology, such as tic location, severity and premonitory urge. Prior studies suggest bilateral abnormalities in the digit representations of individuals with FHD were evident in patients with greater symptom severity and durations of FHD [35–36]. Additionally, repetitive behaviours of the hand, paired with abnormalities in cortical inhibition, were proposed to drive the abnormalities demonstrated in sensorimotor digit representations [37–38]. However, in this study, levels of cortical inhibition, facial tic severity and PU severity were not examined. Consequently, the relationship between individual facial tic severity and sensorimotor representations cannot be determined.

## Conclusion

Sensorimotor mapping of voluntary facial movements do not differ between individuals with TS and TD controls, consistent with evidence of preserved neural activity during voluntary movements. However, unique maps relating to more common tics, such as blinking, may show greater group differences. Additionally, differential SMA activations were evident during volitional movements in the TS group. This could reflect abnormalities within SMA activity influenced by movement complexity, tic expression, or suppression.

## Acknowledgements

CMS was supported by the Precision Imaging Beacon, University of Nottingham and is currently supported by a Postdoctoral Research Fellowship funded by a philanthropic donation from Daniel Katz Ltd. MSH and SRJ were supported by the NIHR Nottingham Biomedical Research Centre (BRC). MA and STF were supported by the Medical Research Council (MRC) (MR/M022722/1). SRJ was supported by the MRC (MR/T032588/1).

Funding from NIHR Nottingham BRC and MRC (MR/T032588/1) supported data collection, participant inconvenience allowances and travel expenses. The views expressed are those of the authors and not necessarily those of the NHS, the NIHR or the MRC.

## Author contributions

**Caitlin M. Smith**: Conceptualisation, Methodology, Formal analysis, Investigation, Data Curation, Writing – Original draft, Visualisation. **Mairi S. Houlgreave**: Investigation, Formal Analysis, Writing – Review & Editing. **Michael Asghar**: Methodology, Formal Analysis, Writing – Review & Editing. **Susan T. Francis**: Conceptualisation, Methodology, Writing – Review & Editing, Supervision. **Stephen R. Jackson**: Conceptualisation, Methodology, Writing – Review & Editing, Supervision, Project administration, Funding acquisition.

## Supplementary information

**Supplementary information 1.**
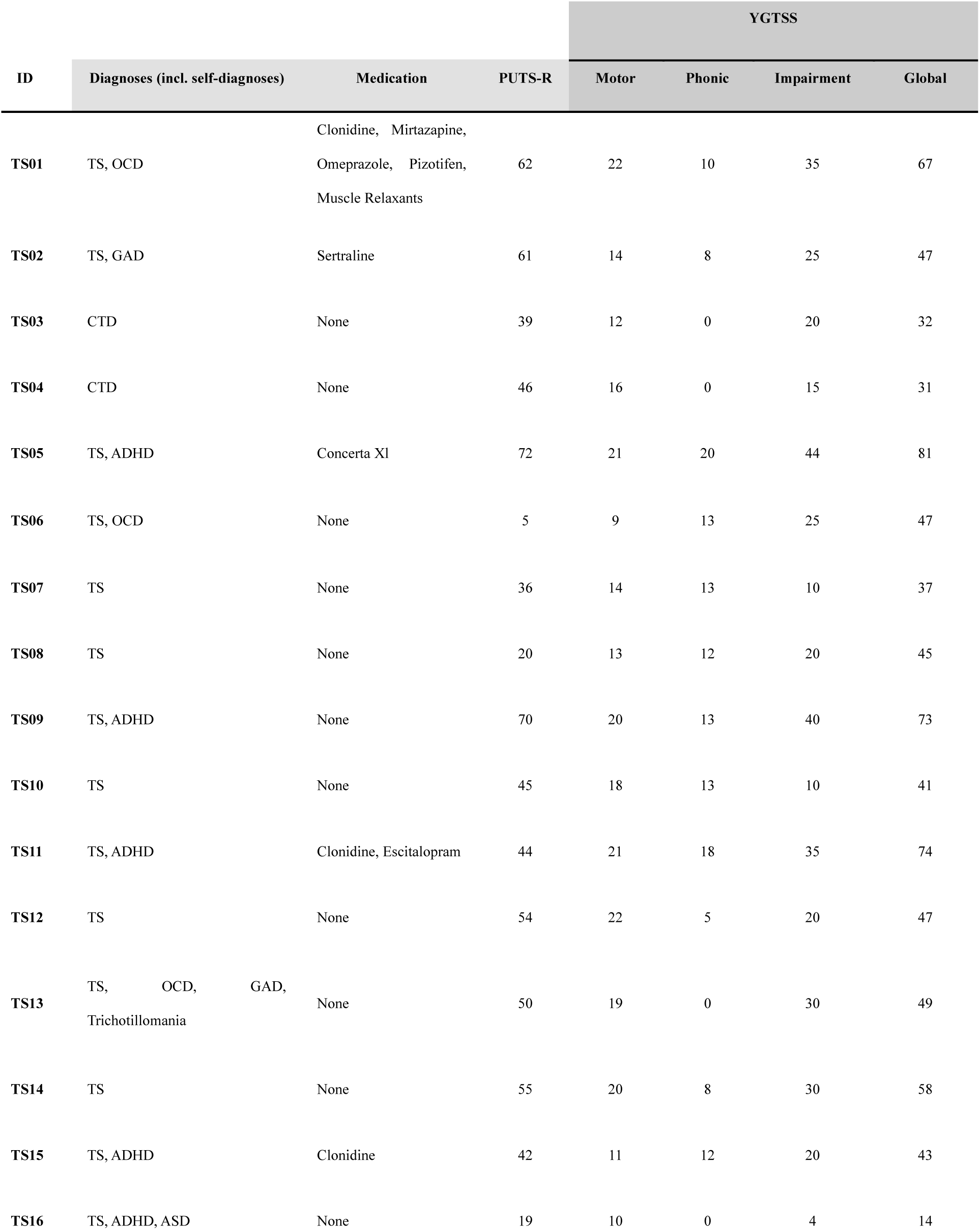
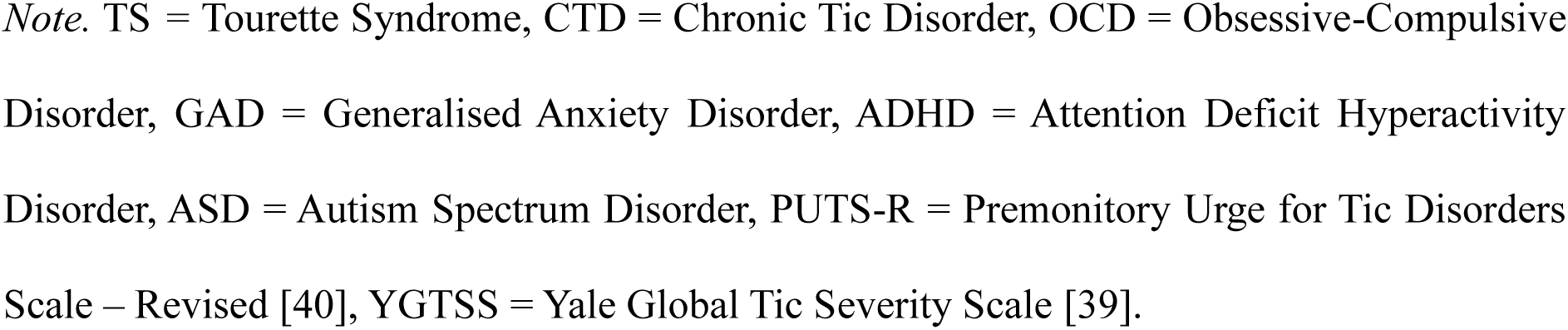
Diagnoses, medication and clinical scores of TS participants.

*Diagnoses, medication and clinical scores of TS participants*
| ID | Diagnoses (incl. self-diagnoses) | Medication | PUTS-R | YGTSS |  |  |  |
| --- | --- | --- | --- | --- | --- | --- | --- |
|  |  |  |  | Motor | Phonic | Impairment | Global |
| TS01 | TS, OCD | Clonidine, Mirtazapine,<br>Omeprazole, Pizotifen,<br>Muscle Relaxants | 62 | 22 | 10 | 35 | 67 |
| TS02 | TS, GAD | Sertraline | 61 | 14 | 8 | 25 | 47 |
| TS03 | CTD | None | 39 | 12 | 0 | 20 | 32 |
| TS04 | CTD | None | 46 | 16 | 0 | 15 | 31 |
| TS05 | TS, ADHD | Concerta XL | 72 | 21 | 20 | 44 | 81 |
| TS06 | TS, OCD | None | 5 | 9 | 13 | 25 | 47 |
| TS07 | TS | None | 36 | 14 | 13 | 10 | 37 |
| TS08 | TS | None | 20 | 13 | 12 | 20 | 45 |
| TS09 | TS, ADHD | None | 70 | 20 | 13 | 40 | 73 |
| TS10 | TS | None | 45 | 18 | 13 | 10 | 41 |
| TS11 | TS, ADHD | Clonidine, Escitalopram | 44 | 21 | 18 | 35 | 74 |
| TS12 | TS | None | 54 | 22 | 5 | 20 | 47 |
| TS13 | TS, OCD, GAD,<br>Trichotillomania | None | 50 | 19 | 0 | 30 | 49 |
| TS14 | TS | None | 55 | 20 | 8 | 30 | 58 |
| TS15 | TS, ADHD | Clonidine | 42 | 11 | 12 | 20 | 43 |
| TS16 | TS, ADHD, ASD | None | 19 | 10 | 0 | 4 | 14 |

*Note.* TS = Tourette Syndrome, CTD = Chronic Tic Disorder, OCD = Obsessive-Compulsive Disorder, GAD = Generalised Anxiety Disorder, ADHD = Attention Deficit Hyperactivity Disorder, ASD = Autism Spectrum Disorder, PUTS-R = Premonitory Urge for Tic Disorders Scale – Revised [40], YGTSS = Yale Global Tic Severity Scale [39].

**Supplementary Information 2.**
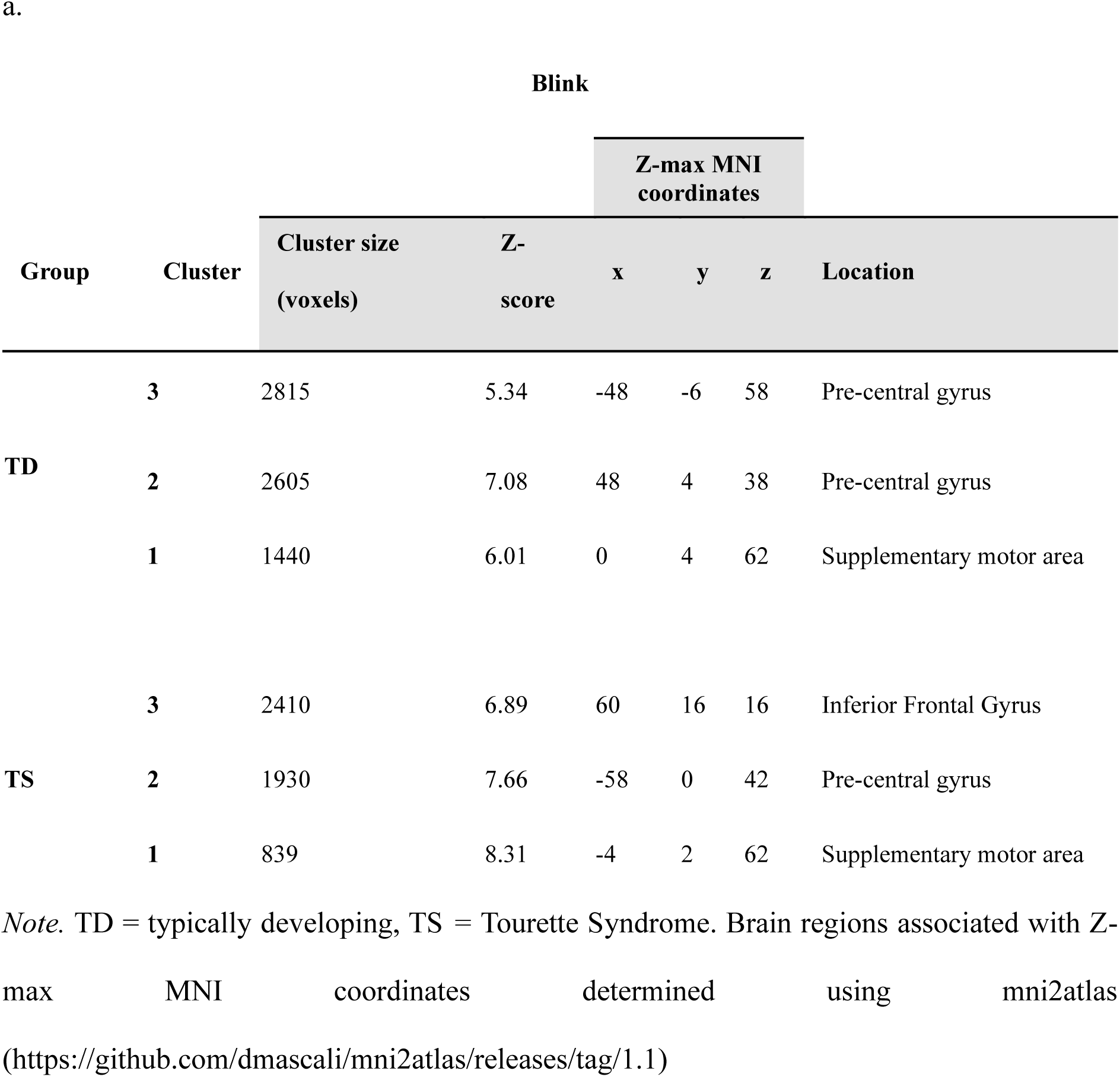
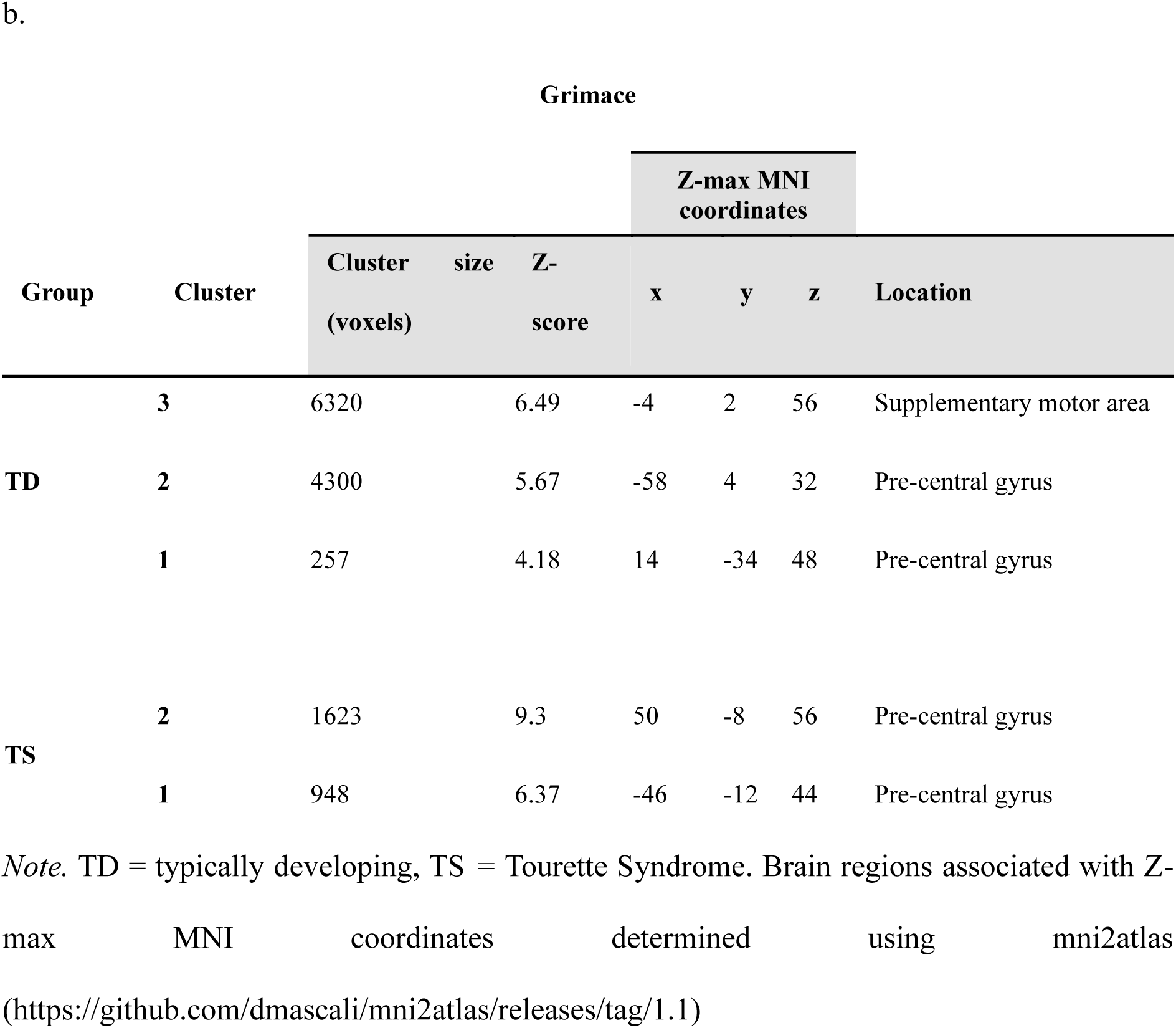
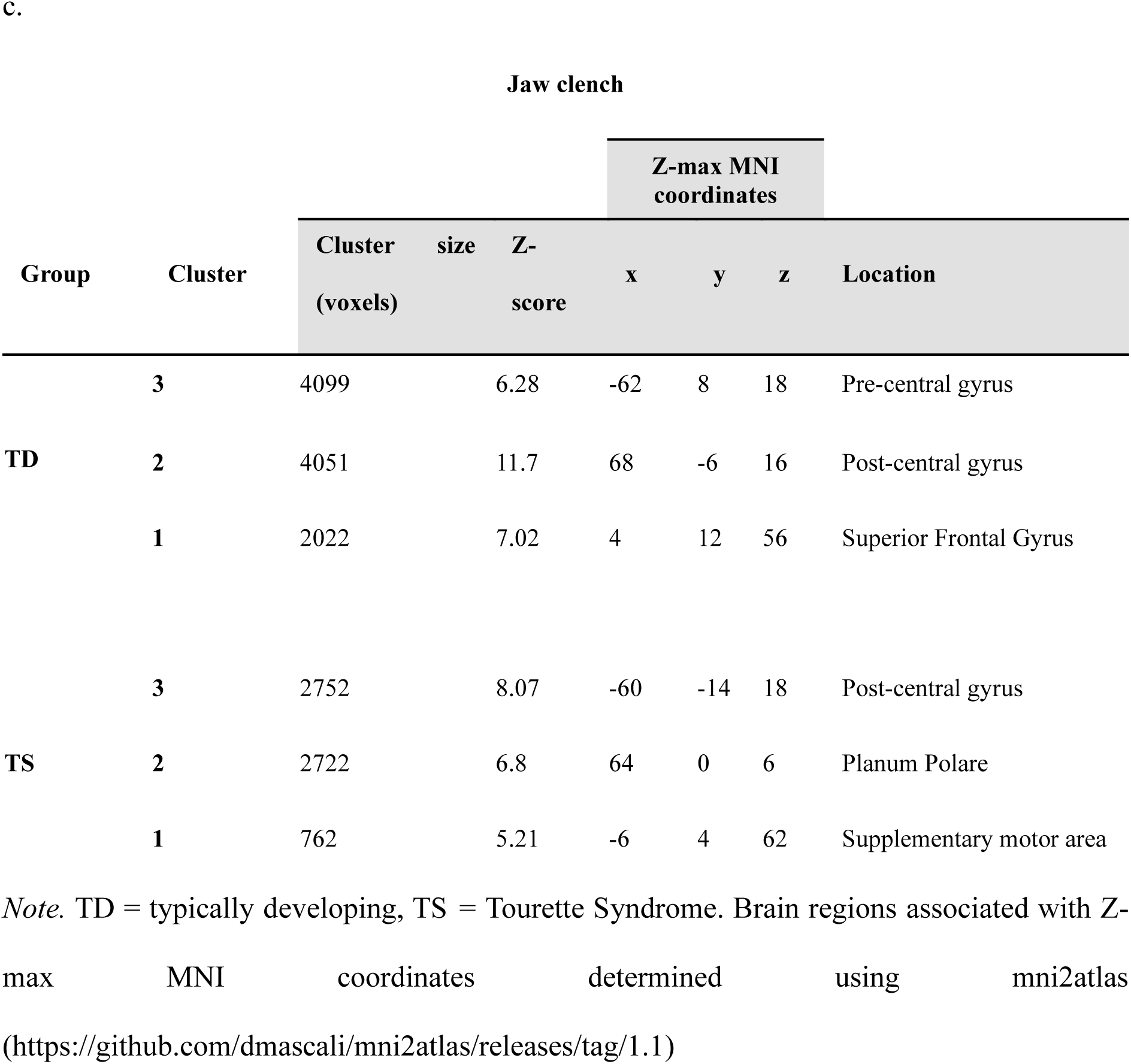
Significant cluster activations for (a) blink, (b) grimace, and (c) jaw clench.

## Notes

### Competing Interest Statement

The authors have declared no competing interest.

